# Response to Shah *et al*: Using high-resolution variant frequencies empowers clinical genome interpretation and enables investigation of genetic architecture

**DOI:** 10.1101/384271

**Authors:** Nicola Whiffin, Angharad Roberts, Eric Minikel, Zach Zappala, Roddy Walsh, Anne H O’Donnell-Luria, Konrad J Karczewski, Steven M Harrison, Kate L Thomson, Helen Sage, Alexander Y Ing, Paul J R Barton, Stuart A Cook, Daniel G MacArthur, James S Ware

## Main text

Recent work by Shah and colleagues^1^ demonstrated that many variants in the ClinVar database^2^ are misclassified, and that disease-specific allele frequency (AF) thresholds can identify wrongly classified alleles by flagging variants that are too prevalent in the population to be causative of rare penetrant disease. While we agree with the main conclusions of this work, the authors compare their AF filtering approach to our recently published method^3^, concluding that the method we advanced “may be prone to removing potentially pathogenic variants”. This is incorrect. Here we demonstrate that our approach is robust, and further illustrate the power of disease-specific AF thresholds for investigating the genetic architecture of disease.

Both methods compare the population frequency of a variant with the prevalence of a disease. However, we advocate considering a fuller definition of disease architecture that explicitly incorporates penetrance and genetic heterogeneity. Using these parameters, we define the maximum AF at which a variant can be observed in the general population to be a credible candidate to cause a defined disease, under the specified genetic architecture. Importantly, we consider each ethnic sub-population separately (‘popmax’), and account for sampling variance in reference datasets^4^. We previously demonstrated that our framework markedly improves signal:noise for identification of penetrant Mendelian variants, without loss of sensitivity^3^.

It is worth restating some important caveats of filtering variants using AF. We must be vigilant for population-specific founder variants, especially in populations where the disease architecture may be distinct, such as the Finnish and Ashkenazi Jewish populations in the Genome Aggregation Database (gnomAD). Also, we do not filter variants observed as singletons due to the stochastic nature of population sampling.

Shah *et al*. report that, in their hands, our method inappropriately filters 15 “high confidence” Pathogenic/Likely Pathogenic ClinVar variants across five cardiac phenotypes (dilated cardiomyopathy, hypertrophic cardiomyopathy, arrhythmogenic right ventricular cardiomyopathy, long QT syndrome (LQT), and Brugada syndrome). The work of Shah *et al*. cannot be directly replicated as their methods and reference dataset are not fully available. Therefore, we have assessed these 15 variants using the larger and more comprehensive gnomAD dataset. We calculated a maximum credible population AF for a variant causative of each disease (defined as in Table 1 of Whiffin *et al*. - see Supplementary Table 1) given a minimum penetrance of 50%.

**Table 1:** Curations and penetrance estimates of 15 variants flagged by Shah *et al*. and 2 additional candidate low-penetrance variants identified in this analysis. Full details can be found in Supplementary Table 3. In the Shah et al analysis 15 variants with “high-confidence” pathogenic assertions in ClinVar were reported to exceed the maximum credible population allele frequency for a pathogenic variant defined by our framework. In our reanalysis, four of these are not filtered by our recommended application of this approach (using gnomAD reference populations), five do not have sufficient evidence for a pathogenic assertion, and six are likely pathogenic but with low penetrance.

| Phenotype | Gene | Cdna | Penetrance | ACMG evidence | ACMG class | conclusion |
| --- | --- | --- | --- | --- | --- | --- |
| HCM | *MYBPC3* | c.3330+5G>C | 12.0% (3.0-45.0%) | PVS1, PS4, PP1_strong | Pathogenic | Likely pathogenic but with low penetrance |
| LQTS | *KCNQ1* | c.1588C>T | 2.5% (0.64-9.8%) | PVS1, PS4_moderate | Likely Pathogenic | Likely pathogenic but with low penetrance |
| LQTS | *KCNQ1* | c.1781G>A | 4.1% (1.0-16%) | PS4, PM1, PP1, PS3_supporting | Likely Pathogenic | Likely pathogenic but with low penetrance |
| Brugada | *SCN5A* | c.3956G>T | 1.1% (0.010-12%) | PS3, PP2, PP3, PP1 | Likely Pathogenic | Likely pathogenic but with low penetrance |
| Brugada | *SCN5A* | c.5129C>T | Ethnicity matched case data not available | PS3, PM1, PP3 | Likely Pathogenic | Likely pathogenic but with low penetrance |
| Brugada/<br>LQTS | *SCN5A* | c.1099C>T | Ethnicity matched case data not available | PS3, PM1, PP3 | Likely Pathogenic | Likely pathogenic but with low penetrance |
| LQTS | *SCN5A* | c.4877G>A | - | PS3_moderate, PP2, PP3 | VUS | Insufficient evidence for a pathogenic assertion |
| LQTS | *KCNQ1* | c.364dupT | - | PVS1 | VUS | Insufficient evidence for a pathogenic assertion |
| HCM | *MYL3* | c.170C>G | - | PS3_moderate, PP1_moderate, PP2 | VUS | Insufficient evidence for a pathogenic assertion |
| LQTS | *SCN5A* | c.5872C>T | - | PVS1 | VUS | Insufficient evidence for a pathogenic assertion |
| LQTS | *KCNQ1* | c.1085A>G | - | PM1, PP3 | VUS | Insufficient evidence for a pathogenic assertion |
| DCM | *LMNA* | c.961C>T | - | PVS1, PP1_Strong, PM2, PS4_Moderate, PS3_Moderate | Pathogenic | Not filtered using gnomAD |
| Brugada | *SCN5A* | c.4885C>T | - | PVS1, PS3_moderate | Likely Pathogenic | Not filtered using gnomAD |
| LQTS | *KCNQ1* | c.573_577delG<br>CGCT | - | PVS1, PP1_supporting, PS3_supporting | Likely Pathogenic | Not filtered using gnomAD |
| HCM | *MYBPC3* | c.3181C>T | - | PVS1, PP1_strong | Pathogenic | Not filtered using gnomAD |
| LQTS | *KCNE1* | c.226G>A | 1.7% (0.58-5.2%) | PS4, PS3 | Pathogenic | Likely pathogenic but with low penetrance* |
| LQTS | *KCNQ1* | c.1664G>A | 1.6% (0.22-12%) | PM5, PM1, PS3_moderate, PP3 | Likely Pathogenic | Likely pathogenic but with low penetrance* |

\*Variants identified by wider gnomAD analysis

With proper application of our approach in this reference population, four (of 15) variants flagged by Shah *et al*. are not filtered (Table 1). We curated the remaining 11 variants according to contemporary ACMG/AMP guidelines^5^ using cardioclassifier.org^6^, ClinVar, and the published literature. Five did not reach a Pathogenic/Likely Pathogenic classification (Table 1; Supplementary Table 3).

The remaining six variants did have sufficient evidence to be classified as (Likely) Pathogenic. For four of the variants, where ethnicity specific case AFs were available, we estimated their penetrance by comparing the case AFs to gnomAD, as previously described^7^. The penetrance of these variants ranged from 1.1% to 12% (Table 1). Crucially, the upper confidence intervals of all six penetrance estimates are well below the pre-specified 50% threshold. In other words, our approach appropriately filters these variants as incompatible with the specified genetic architecture.

We extended our analysis to evaluate all Pathogenic/Likely Pathogenic ClinVar variants for these five cardiac phenotypes. Starting with the same ClinVar VCF (clinvar_20170905.vcf.gz), we annotated variants reported to cause the specified diseases with the tiering strategy outlined by Shah *et al*.^1^ To identify variants above the maximum credible AF for each disease, we used the highest filtering allele frequency across all gnomAD populations (“popmax”) for each variant represented, as described previously^3^. These data, and the code to reproduce this analysis, are available for download from https://github.com/ImperialCardioGenetics/ResponseToShahEtAl.

47 additional variants, previously reported as Pathogenic/Likely Pathogenic, were flagged as “insufficiently rare” by this analysis. We reassessed the clinical interpretation using contemporary ACMG/AMP^5^ guidelines. 45/47 (95.7%) were classified as VUS (Table 2; Supplementary Table 2), and two were classified as (Likely) Pathogenic, but with low penetrance, clearly below our defined genetic architecture (Table 1).

**Table 2:**
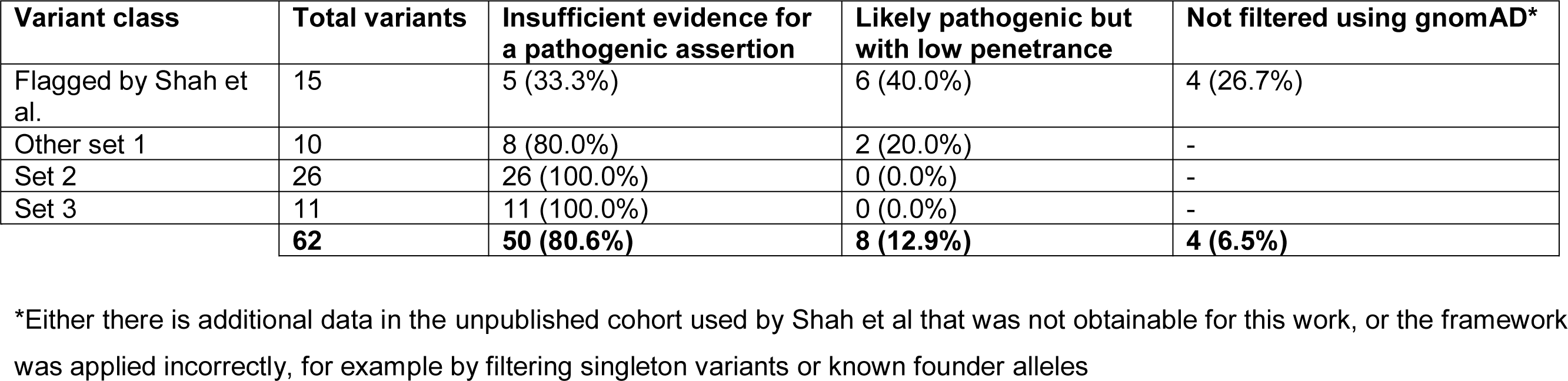
The final classifications of 15 variants flagged by Shah et al. and 47 additional variants identified in this analysis.

| **Variant class** | **Total variants** | **Insufficient evidence for a pathogenic assertion** | **Likely pathogenic but with low penetrance** | **Not filtered using gnomAD\*** |
| --- | --- | --- | --- | --- |
| Flagged by Shah et al. | 15 | 5 (33.3%) | 6 (40.0%) | 4 (26.7%) |
| Other set 1 | 10 | 8 (80.0%) | 2 (20.0%) | - |
| Set 2 | 26 | 26 (100.0%) | 0 (0.0%) | - |
| Set 3 | 11 | 11 (100.0%) | 0 (0.0%) | - |
|  | **62** | **50 (80.6%)** | **8 (12.9%)** | **4 (6.5%)** |

\*Either there is additional data in the unpublished cohort used by Shah et al that was not obtainable for this work, or the framework was applied incorrectly, for example by filtering singleton variants or known founder alleles

Across both analyses, we identified eight (Likely) Pathogenic, low-penetrance variants (Table 2), two of which recapitulate a known mechanism of low-penetrance. These variants are reported as disease-causing for Jervell Lange-Nielsen syndrome (LQT & deafness) in biallelic states, but have low-penetrance for dominant LQT in heterozygous relatives^8^. As these examples demonstrate, our AF filtering framework effectively discriminates alleles with lower penetrance that require tailored counselling.

Within the ACMG/AMP framework, frequency evidence favouring a benign interpretation (BS1) does not preclude a (Likely) Pathogenic classification overall. For example, a common low-penetrance variant may be seen recurrently in cases, showing statistical enrichment in cases over controls (PS4). This contradictory evidence should trigger closer inspection and lead to consideration of a low-penetrance architecture. In other contexts, a more conservative low-penetrance architecture may be specified from the outset.

In conclusion, we previously introduced a statistically-robust, disease-specific framework to leverage reference population AF for variant assessment. We show that this method does not remove true pathogenic variants, provided that they fall within the pre-defined genetic architecture. Although a specified architecture may lead to some low-penetrance variants being flagged with evidence in favour of benign status (BS1), this does not prevent them achieving an actionable ACMG/AMP classification in combination with other lines of evidence. Indeed, flagging this group of variants for in depth review enables more nuanced reporting and counselling around low-penetrance variation.

## Supplemental Data description

Supplementary Data include three tables.

## Acknowledgments

This work was supported by the Wellcome Trust (107469/Z/15/Z), the Medical Research Council (UK), the NIHR Biomedical Research Unit in Cardiovascular Disease at Royal Brompton & Harefield NHS Foundation Trust and Imperial College London, the Fondation Leducq (11 CVD-01), a Health Innovation Challenge Fund award from the Wellcome Trust and Department of Health, UK (HICF-R6–373), and by the National Institute of Diabetes and Digestive and Kidney Diseases and the National Institute of General Medical Sciences, and the National Human Genome Research Institute of the NIH (awards U54DK105566, R01GM104371, and UM1HG008900). E.V.M. is supported by the National Institutes of Health under a Ruth L. Kirschstein National Research Service Award (NRSA) NIH Individual Predoctoral Fellowship (F31) (award AI122592-01A1). A.H.O.-L. is supported by National Institutes of Health under Ruth L. Kirschstein National Research Service Award 4T32GM007748.

This publication includes independent research commissioned by the Health Innovation Challenge Fund (HICF), a parallel funding partnership between the Department of Health and the Wellcome Trust. The views expressed in this work are those of the authors and not necessarily those of the Department of Health or the Wellcome Trust.

## Web Resources

Github repository: https://github.com/ImperialCardioGenetics/ResponseToShahEtAl

